# *SurEau*.c : a mechanistic model of plant water relations under extreme drought

**DOI:** 10.1101/2020.05.10.086678

**Authors:** Hervé Cochard, François Pimont, Julien Ruffault, Nicolas Martin-StPaul

## Abstract

We describe the operating principle of the detailed version of the soil-plant-atmosphere model ***SurEau*** that allows, among other things, to predict the risk of hydraulic failure under extreme drought. It is based on the formalization of key physiological processes of plant response to water stress. The hydraulic functioning of the plant is at the core of this model, which focuses on both water flows and water pools using variable hydraulic conductances. The model considers the elementary flow of water from the soil to the atmosphere through different plant organs (roots, trunk branches, leaves and buds) that are described by their symplasm and their apoplasm compartments. Within each organ the flow of water between the apoplasm and the symplasm is also represented; as well as the flow outside the system, from the symplasm of each organ to the atmosphere, through the cuticular conductance. For each organ, the symplasm is described by a pressure volume curves and the apoplasm by the vulnerability curve to cavitation of the xylem. The model can thus compute the loss of conductance caused by cavitation, a leading mechanisms of plant desiccation and drought-induced mortality. Some example simulations are shown to illustrate how the model works.

## Introduction

Numerous models have been developed to simulate the water relations and gas exchanges of plants under optimal or limiting hydric conditions. These models are based either on empirical relationships or on more mechanistic bases, i.e., based on a physical representation of the physiological processes. A few years ago, we identified that there was no mechanistic model taking into account the water relationships of plants under conditions of extreme water stress, i.e. when the plant reaches its survival limit. This is the reason why we began developing such a model in 2015, first in a simplified form in an Excel spreadsheet, then as a R script (Martin-StPaul et al 2017). These earlier versions used a quasi-static approach and the plant was reduced to two compartments (one symplasmic and one apoplasmic). From 2017 onwards, we developed a new dynamic version of this model in C language, based on a plant segmentation in different organs. This model has already been used in a number of recent publications (Martin-StPaul et al 2017; Duursma et al 2019, Scoffoni et al 2018, Cochard 2019, Brodribb et al 2019, Brodribb et al 2020, Dayet et al 2020, Lamarque et al 2020), but never formally described as here.

The innovative aspect of this model is to describe the temporal evolution of a plant’s water status beyond the point of stomatal closure. Under these extreme stress conditions, the model describes the residual transpiration flow through the cuticle, cavitation processes, and the solicitation of the plant’s water reservoirs. Hence the model also allows to track water quantities in the different plant organs. The objective was also to model these processes both for plants under controlled conditions, as well as under natural current and future conditions.

The ***SurEau*** model is primarily an hydraulic model computing water flows. It can be combined with a simplified photosynthesis, energy budget and growth modules, but those are not described here.

The soil-plant-atmosphere system is segmented and described using different linked hydraulic organ compartments exchanging water fluxes called computational cells, or simply “cells” (Figure 1). These fluxes are determined by gradients of water potential between cells and hydraulic conductances of these cells. The water quantity of each cell is therefore described as a result of incoming and outgoing fluxes; and the water potential of each cell is computed with the appropriate formulation according to the nature of these cells (soil, symplasm, apoplasm): (i) a pedo-transfer function for the soil (Van Genuchten et al 1980); (ii) a pressure-volume curves for the symplasm (Tyree and Hammel 1972), which expresses the relationship between water content and water potential and (iii) a vulnerability curve to cavitation and the capacitance in the case of the apoplasm (Cruiziat et al 2002).

**Figure 1:**
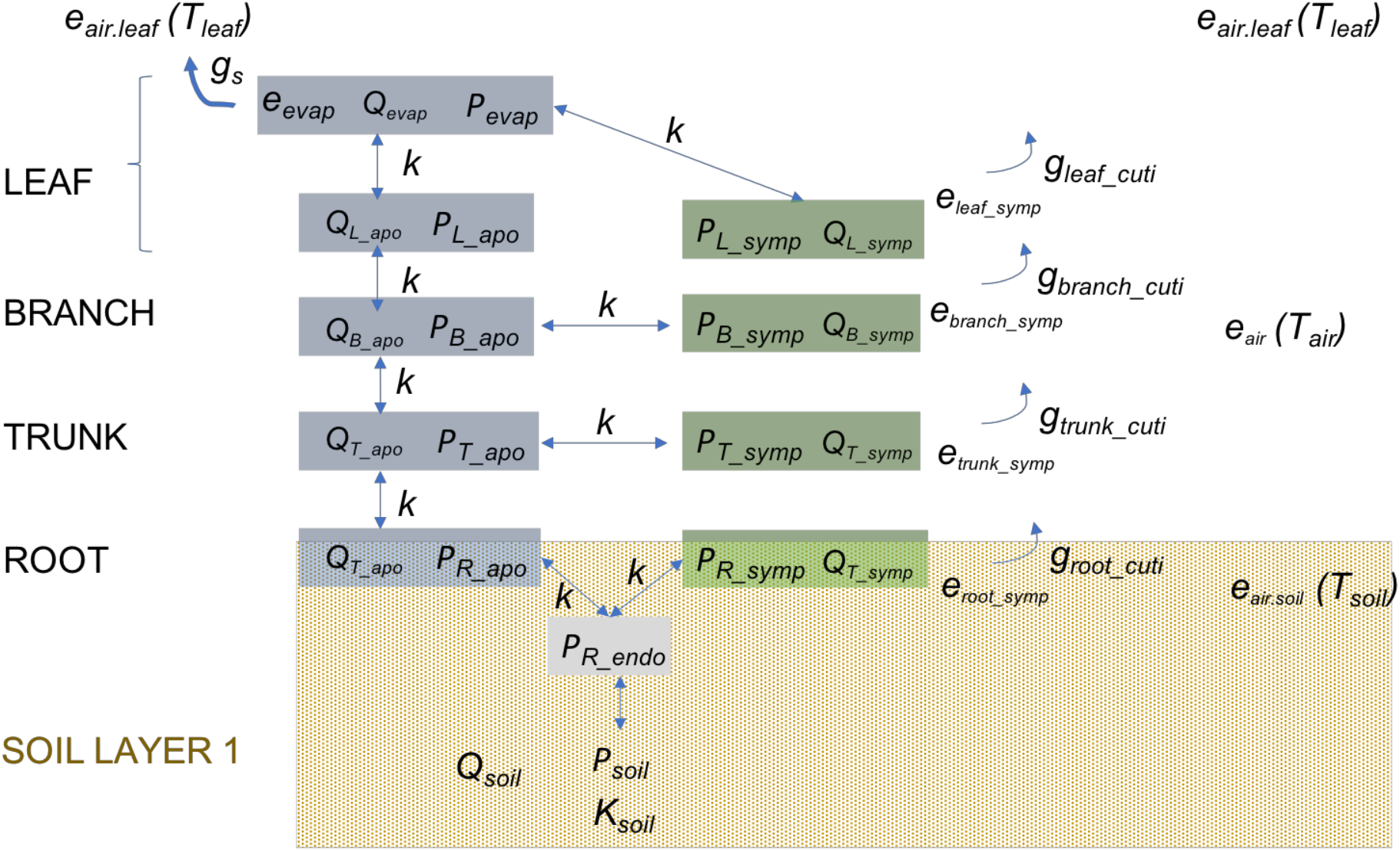
Simplified representation of the architecture of the ***SurEau***.c model in relation to the environment. *Q* and *P* the water quantity and water potential defined within a compartment (or “computational cell”), *K* the hydraulic conductance defined between two compartments (materialized by arrows), *g* the gas phase conductance, *e* the actual vapour pressure and *T* the temperature.

In order to model explicitly the dynamics of the system, the model is integrated over a very small time step (*dt*), on the order of milliseconds, to avoid numerical instabilities associated with the Courant-Friedrichs-Lewy condition (CFL, Dutykh 2016). Here we provide a full description of the model. This is not a user guide, which will be released in the future together with the ***SurEau*** code. An object-oriented version embed into the *Capsis* platform (Dufour-Kowalski et al 2012) is also under development: http://capsis.cirad.fr/capsis/help_en/sureau

## I Formalization of the soil-plant-atmosphere system in *SurEau* (Figure 2)

The soil-plant-atmosphere continuum is idealized as a collection of five linked organ types (roots, trunk, branches, buds, leaves), each of them containing both an apoplasmic and a symplasmic compartment. In addition, roots also have an endoderm and leaves also have an evaporative site. Each compartment is a computational cell of the model. The root system is divided into 3 elements, each occupying a different soil horizon and being connected to the trunk. The crown of the tree is divided into *n* identical branches connected from the trunk in parallel. Therefore, the number of branches has no incidence on the water transport to the canopy that can still be treated using the “big leaf’ approximation, unless some climate variability is considered within the crown.

**Figure 2:**
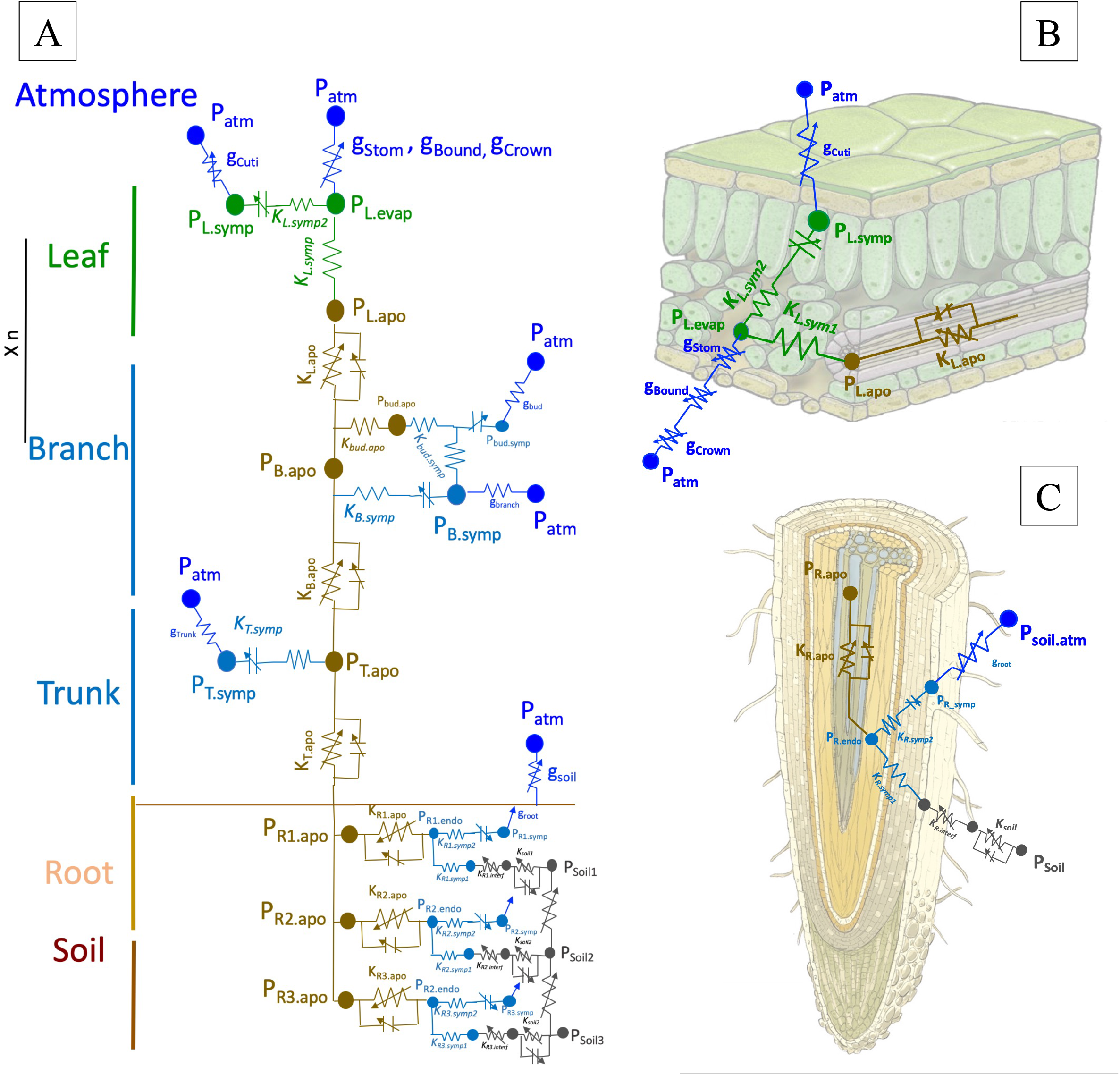
Idealization of the soil-plant-atmosphere continuum in *Sureau*.c. The plant is described as a network of conductances and capacitances. In A) is represented the whole architecture of the model. B) and C) show the formalizations for leaves and roots, respectively.

The state variables are presented in Table 1. They include the water potentials (*P*, but also osmotic potential *π*, and turgor pressure, *Tp*) and the water quantity (*Q*). Capacitances (C) and conductance (*K*) are parameters that can vary with temperature or when the cavitation occurs in xylem vessels or when leaf fall occurs (Table 2). Hence the percent loss of conductance (*PLC*) and the percent leaf fall (*PLF*) are also considered as state variables (Table 1). A simplified representation of the system is presented in Figure 1 and a more detailed representation in Figure 2. In what follows, indices are used to designate the different compartments or organs. For organs, the following index is used: L for *leaf*, B for *branches*, bud for *bud*, T for *trunk* and R for *roots*. As there may be several roots or branches in parallel, we also use the index 1, 2,… n to designate the organ/number/soil layer. For the compartments, the indices *apo* or *symp* are used to designate the *apoplasm* or *symplasm* respectively. For roots, there is an additional compartment, the *endoderm*, which is designated by the index *endo*. A point separates the indices of the organ and the compartment. Thus, for example, the water potential of the leaf apoplasm is written *P_L.apo_*.

**Table 1:** State variables (that characterize the state of the soil-plant system) used in ***SurEau***.

| State variables | Description | Unit | Type of “cell” |
| --- | --- | --- | --- |
| $Q$ | Water quantity | mmol | *apoplasm, symplasm* |
| $P$ | Water potential, | MPa | *apoplasm, symplasm* |
| $\pi$ | Osmotic potential | MPa | *symplasm* |
| $Tp$ | Turgor pressure | MPa | *symplasm* |
| PLC | Percent loss of conductance | % | *apoplasm* |
| PLF | Percent leaf fall | % | - |

**Table 2:**
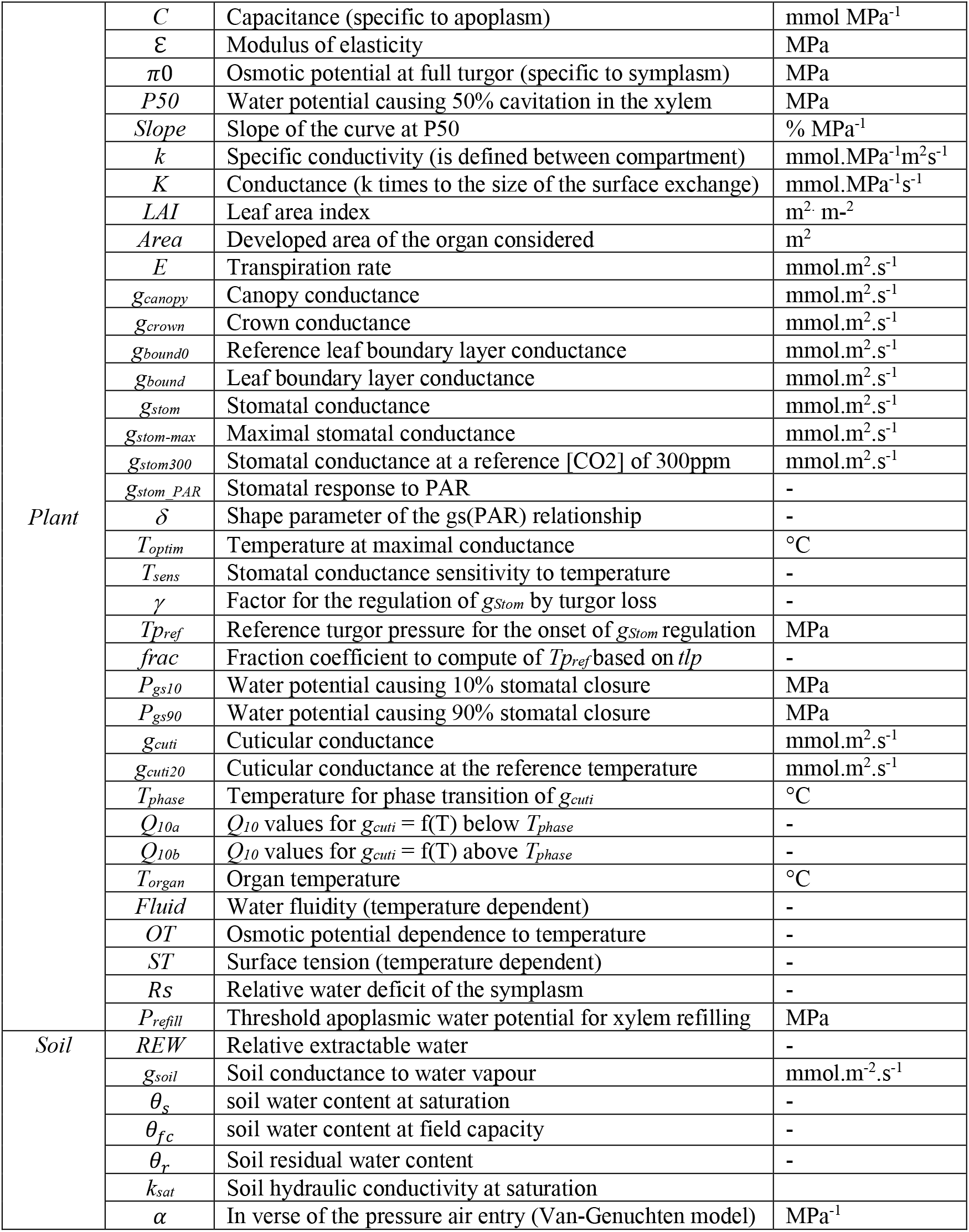
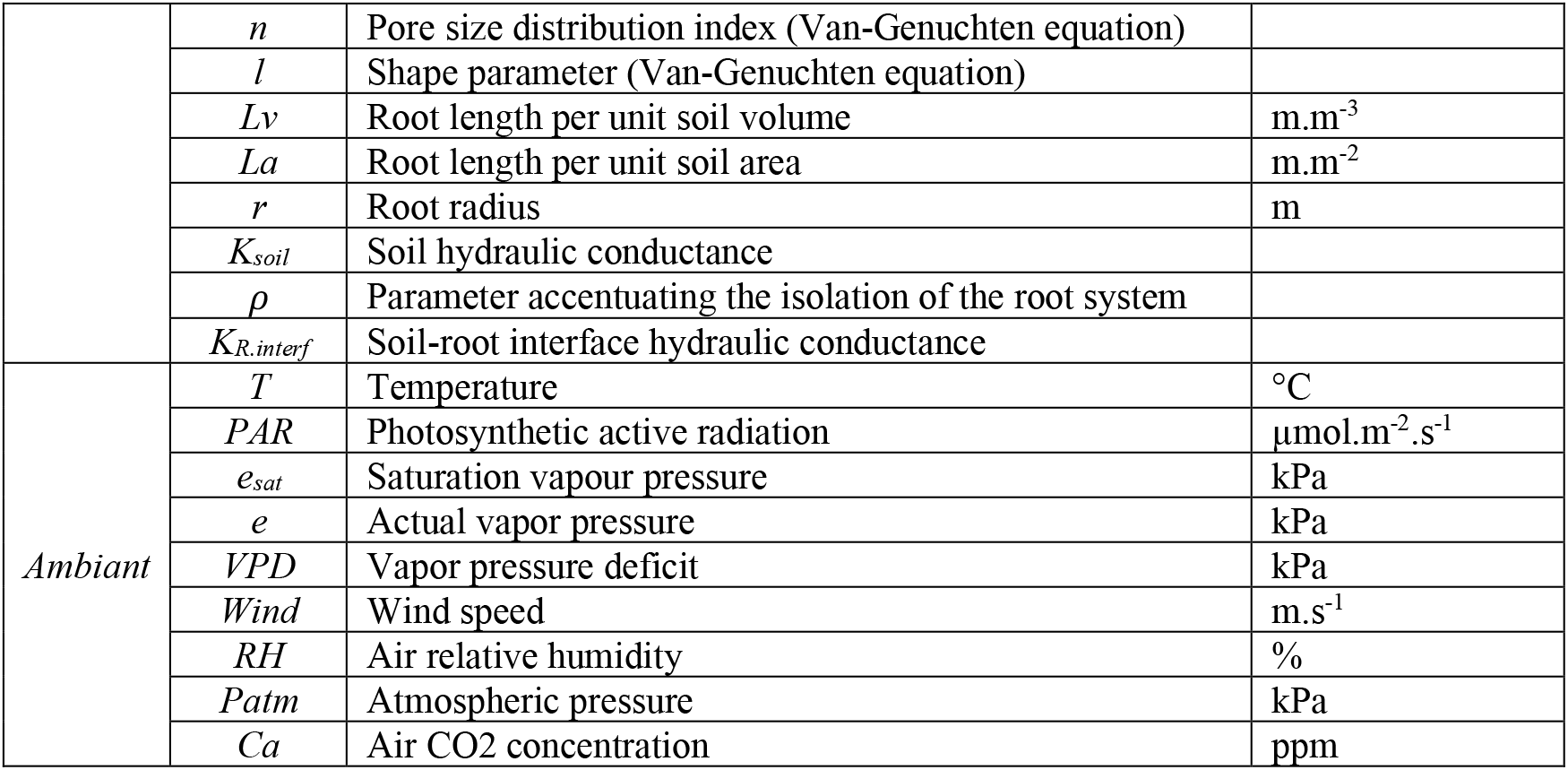
Nomenclature (model variables and parameters used in this document)

Specificities of the water flows in the leaves are shown in Figure 2c. Sap at pressure *P_L.apo_* enters the leaf through the apoplasm conductance (petioles + veins, *K_L.apo_*), then passes through mesophyll cells (*K_L.symp1_*) to reach the site of evaporation in the substomatic chamber (*P_L.evapo_*). This evaporation site is considered as an apoplasmic compartment. The water reservoir of the leaf symplasm is connected through another symplasmic conductance (*K_L.symp2_*) to the evaporation site. This reservoir also directly loses water via cuticular transpiration. For the roots, we consider a formalism quite similar to leaves (Figure 2d). For each root, water from the soil reservoir at *P_soil_* passes through the soil conductance (*K_soil_*), the soil-root interface *K_R.inter_* and the cortical symplasmic layer of the absorbent roots *K_R.symp1_* to reach the endoderm (*P_R.endo_*) and then the apoplasm of the root stele (*K_R.apo_* to *P_R.apo_*). The root symplasmic water reservoir is connected to the endoderm by a conductance *K_R.symp2_* which can also lose water by evaporation through the root periderm.

In general, we preferred to use *mol* as a unit for water movements (instead of *g* or *m*^3^) for consistency with the gaseous water flows through the stomata or cuticle. The list of all parameters and variables used in this document is presented in Table 2.

## II Implementation of *SurEau*.c

The general principles of calculating flows and potentials in ***SurEau*** are the following:

1. Differences in water potentials between compartments create elementary water movement of water molecules (*dq*) at the different interfaces according to fluxes computed with Fick’s using the interface conductance.
2. The water content of a compartment is increased by *incoming* fluxes and lowered by *outgoing* fluxes and transpiration, because of water mass conservation law.
3. The water potential of each apoplasm compartment is derived from the water quantity and capacitance; the water potential of each symplasm compartment is derived from its pressure-volume curve.
4. Transpirations are computed from 1) the vapor pressure deficit at the level of each compartment, and 2) the gas phase conductance which includes the cuticle and the stomatal conductance which accounts for various regulation.

Overall, the model is implemented within two loops:

1. A first loop at a very small time step (*dt* ~ 0.01 s) computing sequentially *dq* (between the different organs and compartments), *Q, P* (and *Pi* and *Tp*), and *K*.
2. To reduce time consumption, an external second loop computes processes that are not affected by numerical instabilities and longer to compute on a larger time step, i.e. with exponential or power functions. It operates on the time scale of seconds to minutes and includes:

- Organ transpiration and temperature. Transpiration and temperature are computed at the leaf surface by using stomatal conductance, leaf cuticular conductance and solving the energy budget. For all compartments (bud, branch, trunk, root, soil) only transpiration is computed using cuticular conductances and by assuming *T_compartment_* = *T_air_* or = *T_soil_*.
- Cavitation and redistribution of water released by cavitation within other compartments.
- The dependency of some physical properties of water solutions (fluidity, surface tension, osmotic potential) to temperature
- Additional processes (photosynthesis, respiration, growth, leaf rain interception etc.) which are not described here.

### a) Small time step 1: computation at *dt* (~0.01 s) of *dq, Q, P, K*

#### 1 Exchanges of water molecules between organs and compartments (*dq*, mmol) during small time steps

These exchanges during *dt* are computed from Fick’s law (with *K* and *P*) that allows to compute the water fluxes between compartments. The description is given below for all compartments in the order they are computed in the model.

<u>Leaf</u>

- Water movement from the leaf apoplasm to the site of evaporation:

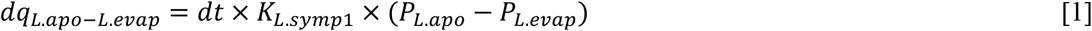
- Water movement from the site of evaporation to the symplasm:

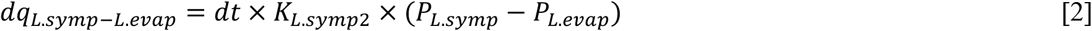

<u>Branch</u>:

- Water movement from the branch apoplasm to the leaf apoplasm:

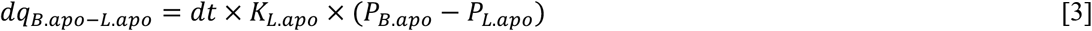
- Water movement from the branch apoplasm to the branch symplasm:

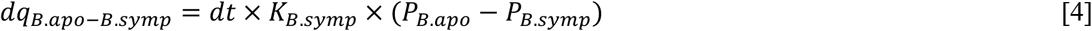

<u>Trunk</u>

- Water movement from the trunk apoplasm to the branch apoplasm:

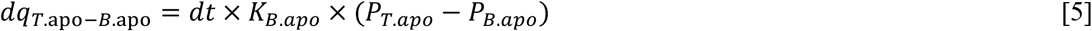
- Water movement from the trunk apoplasm to the trunk symplasm:

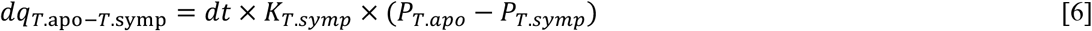

<u>Roots (for root 1 only)</u>

- Water movement from the root apoplasm to the trunk apoplasm:

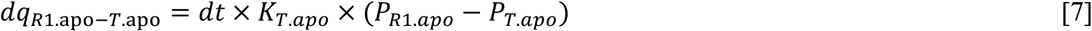
- Water movement from the root endoderm to the root apoplasm:

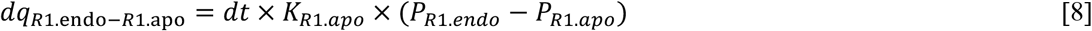
- Water movement from the root symplasm to the root endoderm:

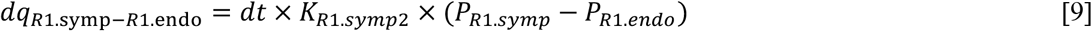
- Water movement from the soil to the root endoderm:

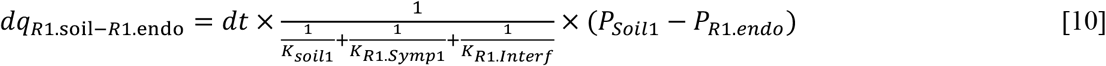

#### 2 Integration over time of the Water quantity (*Q*, mmol)

The integration over small time steps of *Q* is based on the water mass conservation law following a first order explicit scheme. It is described below for all compartments as computed in the model. For each compartment interacting with the atmosphere (e.g. evaporative site, symplasm, etc), sink terms corresponding to transpirations (*dq_Stom_,dq_Cuti_,etc*) are derived from the corresponding gas phase conductances. These terms are computed in loop 2 (part b.1).

<u>Leaf</u>

- Water content of the site of evaporation:

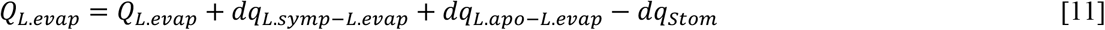

with *dq_Stom_* the water transpired through the stomata, computed from the stomatal conductance (*g_stom_*) and VPD in loop 2.
- Water content of the leaf symplasm:

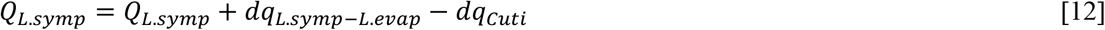

with *dq_Cuti_* the water transpired through the leaf cuticle computed from the gas phase *g_Cuti_* and VPD in loop 2.
- Water content of the leaf apoplasm:

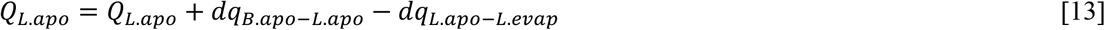

<u>Branch</u>

- Water content of the branch symplasm:

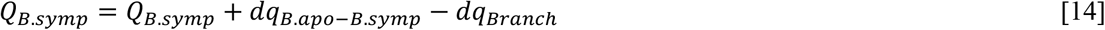

with *dq_Branch_* the water transpired through the branch periderm computed from the gas phase conductance *g_Branch_* and VPD in loop 2.
- Water content of the branch apoplasm:

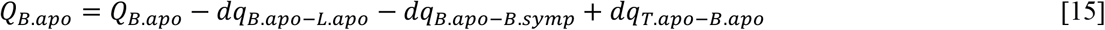

<u>Trunk</u>

- Water content of the trunk symplasm:

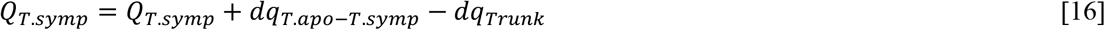

with *dq_Trunk_* the water transpired through the trunk periderm computed from *g_Trunk_* and VPD in loop 2.
- Water content of the trunk apoplasm:

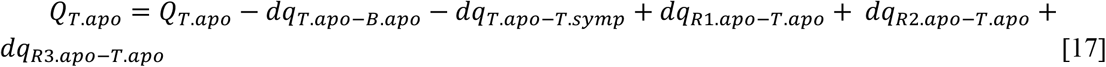

<u>Root</u> (shown for root 1 only)

- Water content of the root symplasm:

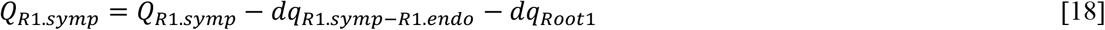

with *dq_Root1_* the water transpired through the root periderm and computed from the conductance *g_Root1_* and VPD in loop 2.
- Water content of the trunk apoplasm:

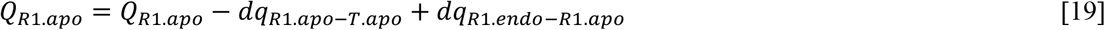
- Water content of the endoderm:

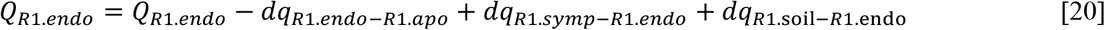 Similar equations are applied to the second and third root elements.

<u>Soil (here only for one « root layer » over three)</u>

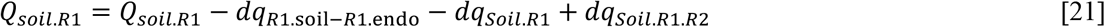

with *Q_soil.R1_* soil water content; *dq_Soil.R1_* soil evaporation, *dq_R1.soil-R1.endo_* water transfer between soil and root endoderm, *dq_Soil.R1.R2_* capillary transfer between soil layers.

The same applies for soil layers two and three.

#### 3 Water potential (*P*, MPa), osmotic potential (*π*, MPa) and turgor pressure (*Tp*, MPa)

They are computed from the variation in water content, i.e., the difference between current water content and its value at full saturation (noted with the subscript 0) and organ traits (*C* the capacitance, ε modulus of elasticity and *π*0 the osmotic potential at full turgor). Here equations are given for the leaf (evaporation site, apoplasm and symplasm). Similar equations apply for branch, trunk and roots.

<u>For the *apoplasmic* water potential</u>

By definition of the capacitance parameters that are assumed constant, we can compute:

- Water potential of the leaf evaporation site:

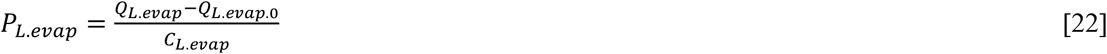
- Water potential of the leaf apoplasm:

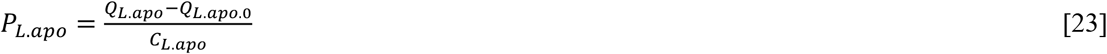

<u>For the *symplasmic* water potential</u>

The symplasmic water potential is derived from the pressure volume equations (Tyree and Hammel 1972), it is subsequently defined as the sum of the turgor pressure *Tp* and osmotic potential *π*

- Water potential of the leaf symplasm:

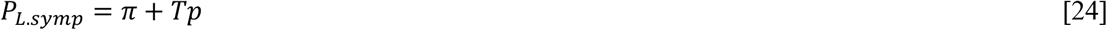 The turgor pressure is computed as

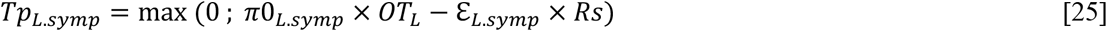

with *π*0 the osmotic potential at full turgor, ε the modulus of elasticity, *OT_L_* is a factor correcting for the effect of temperature on osmotic potential. *OT_L_* is computed at a longer time step, in the second loop. *Rs* is the relative water deficit of the organ symplasm.

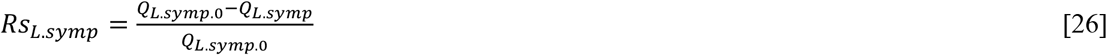 And the osmotic potential (*π*) is computed as:

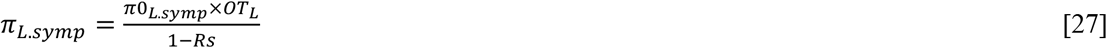 The same equations apply for the symplasm of other organs. For tall trees, the model also accounts for the gravimetric potential due to height.

#### 4 Hydraulic conductances (*K*, mmol s^-1^ MPa^-1^)

Xylem conductances (apoplasmic) vary from their initial value with the degree of cavitation (expressed by the *PLC*, the percent loss in conductivity) and the fluidity of water, which is temperature dependent. In addition to leaf fall and temperature, symplasmic conductances can also depend on other factors, such as, for instance, the effect of aquaporins regulation. Such additional effects are not described here. Leaf fall (if it occurs) also modifies the leaf conductances. The equation is given here for leaves but the same applies for all organs.

- Conductance of the leaf apoplasm:

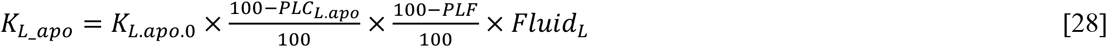
- Conductance of the leaf symplasm:

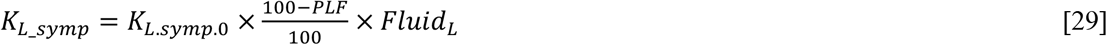

with the *K_L.apo.0_* the initial conductance for the leaf apoplasmic compartment; *PLC* the percent loss of conductivity computed from cavitation, *PLF* the percent leaf fall that is empirically derived, and *Fluid_L._*. the water fluidity computed in loop 2.

### b) Large time step loop *(1 second or minute)*

This loop computes on a larger time step (*dt*100*, or more) processes that are involved in the update of parameter values and boundary conditions of the system. These computations can be time consuming and are not involved in the constraints associated with the CFL condition, which only deals with water fluxes and time integration of water quantities. It includes transpiration (and energy balance of the leaf), cavitation, and the temperature dependence of some physical properties of water (fluidity, surface tension, osmotic potential). Other processes can be computed in this loop (e.g. photosynthesis, growth) but they do not really interact with hydraulics at the time step considered here, so they are not described here.

#### 1 Transpirations (*E*, mmol s^-1^ m^-2^)

The plant loses water through its stomata and its cuticles. The total plant transpiration *E_Plant_* can be decomposed as:

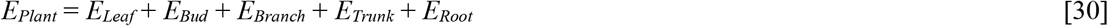

With *E_Leaf_* further decomposed as:

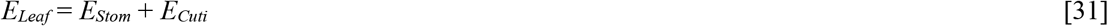

Further including the soil water loss *E_soil_* we can compute the “ecosystem” evapotranspiration *E_Eco_* as:

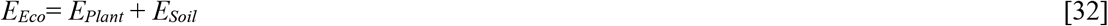

<u>Transpiration computation general principles</u>

The transpiration *E_organ_* (mmol s^-1^ m^-2^) is computed with the gas phase conductance *g_organ_* (mmol s^-1^ m^-2^), the vapor pressure deficit between the organ and the atmosphere *VPD_organ_* (kPa) and the atmospheric pressure *P_atm_* (kPa):

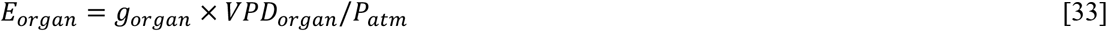

*g_organ_* is constant for all organs, except for the cuticle and the stomata, based on specificities described in the next subsections.

The vapor pressure deficit between the air and organ surface is given by (Cochard 2019):

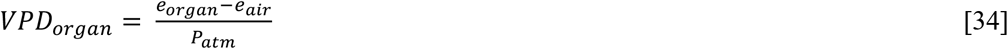

where *e_air_* is the vapor pressure of bulk air and *e_organ_* is the vapor pressure at the level of the organ symplasm. Both are a function of temperature according to Buck’s equation:

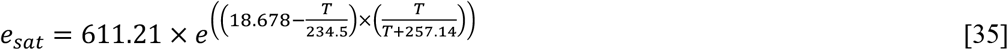

where *T* represents either the organ (*e_sat_organ_*) or the air (*e_sat_air_*) temperature (°C). The leaf temperature is computed from an energy budget model adapted from Sinoquet et al (2001). The branch, bud and trunk temperatures are assumed equal to the air temperature. The root temperature is also equal to the soil temperature.

The actual vapor pressure at the level of the compartment under consideration depends of *e_sat_organ_* and its water status (*P_organ_*) according to:

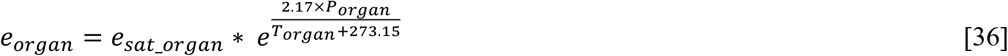

Similarly, the air vapor pressure is a function of air relative humidity *RH*(%):

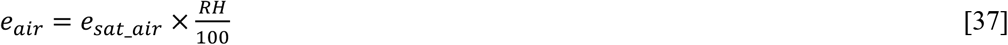

For root evaporation rates, we compute *e_air-soil_* with the soil water potentials.

<u>Transpiration specificities of the leaf interface (stomata and cuticle):</u>

The conductance of the leaf interface with the atmosphere *g_Canopy_* is variable and is composed of four conductances:

- *g_Stom_*: the conductance of the stomatal pores
- *g_Cuti_*: the conductance of the leaf cuticle

these two conductances are in parallel and in series with:

- *g_Bound_*: the conductance of leaf boundary layer
- *g_Crown_*: the aerodynamic conductance of the tree crown.

The canopy foliage conductance *g_Canopy_* is the total conductance given by:

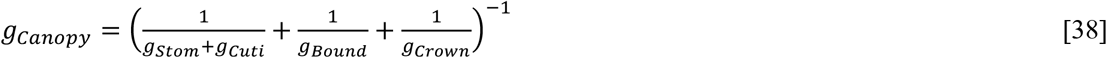

*g_Bound_*, the conductance of the leaf boundary layer is computed following Jones (2013) and varies with leaf shape, leaf size and wind speed.
*g_Crown_*, the conductance of the tree crown varies only with the wind speed (Jones 2013):

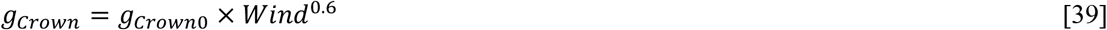

with *g_Crown0_* the reference conductance and *Wind* the wind speed (m.s^-1^).

The conductance of the leaf cuticle *g_Cuti_* is a function of the leaf temperature which is based on a single or double *Q_10_* equation depending whether *T_leaf_* is above or below the transition phase temperature *T_phase_* (Cochard 2019):

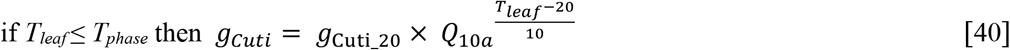

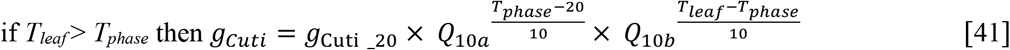

where *g_Cuti_20_* is the leaf cuticular conductance at 20°C and *Q_10a_* and *Q_10b_* are the *Q_10_* values of the relationship below and above *T_phase_*, respectively.

Once *g_Cuti_* is determined, the cuticular transpiration rate *E_Cuti_* can be computed as:

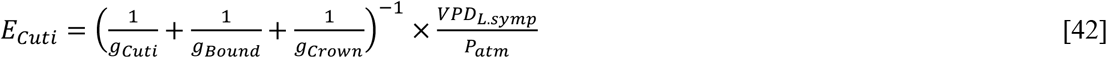

where *VPD_L.symp_* is computed with a formulation similar to other organs (see above)

Stomatal transpiration is a critical part of the ***SurEau*** model. It is based on the regulation of the stomatal conductance. Several options were implemented to compute this part in the next subsection.

<u>Stomatal transpiration</u>:

Stomatal conductance *g_Stom_* is known to respond to multiple variable with the most limiting one determining the actual conductance (Jarvis, 1976). ***SurEau*** takes into account the dependence of *g_stom_* to light, temperature and CO2 concentration on the one hand, and water status on the other. Water status effects can be considered through different representations (*finalist, mechanistic* or *empiric*) that depend on options chosen by the operator and are described in details below.

First, a maximal stomatal conductance *g_Stom_max_* is defined as the minimum value of *g_stom_* dependence to air temperature (*g_Stom_T__*) and to atmospheric CO2 (*g_Stom_CO2_*):

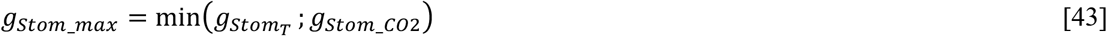

*g_Stom_T__* follows a bell-shape temperature response, parametrized with a maximal value at *T_optim_* and a sensitivity response to temperature *T_sens_*:

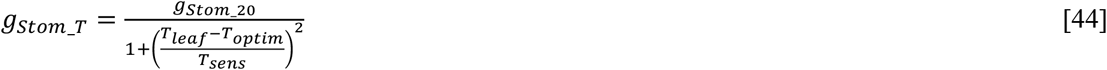

where *g_Stom_20_* is the maximal conductance at 20°C.

*g_Stom_CO2_* depends on atmospheric CO2 concentration Ca (ppm) and is inversely proportional to *Ca*:

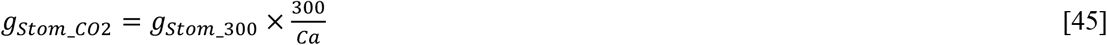

with *g_Stom_300_* the *g_Stom_max_* value at 300 ppm. Once *g_Stom_max_* determined, we use a Jarvis’s like approach to account for both the effect of incident PAR (μmol m^-2^ s^-1^) and the effect of water deficit. First, we introduce *g_Stom_min_*, the minimum *g_Stom_* value when PAR=0 (Duursma et al 2019). We then compute *g_Stom_PAR_* as:

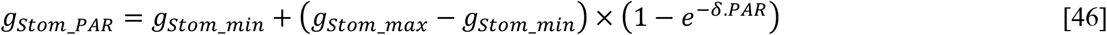

with *δ* a shape parameter.

Knowing *g_Stom_PAR_* we can account for the role of water status for which we implemented three different options corresponding to different vision about how stomata works (*mechanistic, empiric* or *finalist*):

i. A first *mechanistic* option assumes that stomata respond to bulk leaf turgor loss. That is, when the turgor pressure reaches a predefined threshold stomata close. In this case, stomatal conductance respond to changes in leaf turgor pressure and is proportional to *g_Stom_PAR_* by a factor *γ* defined as:

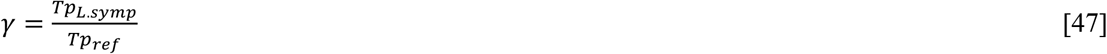

and

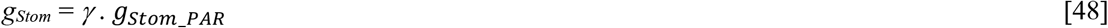

where *Tp_ref_* is a reference turgor pressure that is set equal to the leaf turgor pressure measured at midday on sunny and well-watered conditions. Alternatively, *Tp_ref_* can been set at a fraction (*frac*) of leaf osmotic potential at full turgor (*π*0_*L.symp*_/*frac*) to represent the observation that water turgor pressure affect stomatal conductance only below a certain threshold. A major advantage of this approach is that the stomatal model can be parameterized using leaf pressure volume curves equations available for many species (Bartlett et al 2012; Martin StPaul et al 2017).
ii. A second option uses *empirical* responses of stomata to a proxy of water status (such as leaf water potential (symplasmic) or soil water potential). These can be easily implemented when the soil or leaf symplasmic water potentials corresponding to 0.9*g_Stom_max_*(*P_gs10_*) and 0.1*g_Stom_max_*(*P*_gs90_) are known. In this case, *g_Stom_* is computed as:

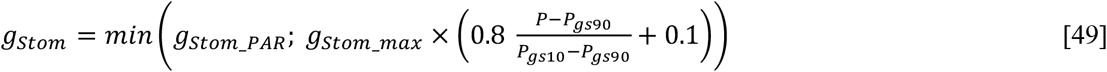 With *P* corresponding to the soil or symplasmic water potential. This also presents the advantage to be easily implemented for multiple species thanks to available databases (*i. e*. the ***SurEau*** database, Martin-StPaul et al. 2017).
iii. A third option is called *finalist*, because *E_Stom_* is regulated assuming that stomata close so that a target organ water potential (e.g. *P_B.apo_*) remains above a threshold potential corresponding to *E_Stom_* = 0. As proposed in (Martin-StPaul et al 2017), this option can be used, for instance, to close stomata at incipient embolism formation in a given compartment (e.g. *P_B.apo.12_*, the branch apoplasmic potential corresponding to a PLC of 12%, Martin-StPaul et al 2017). This regulation requires therefore to first calculate the canopy foliage transpiration (*E_clim_* (mmol s^-1^ m^-2^)) in order to derive the target *P*. *E_clim_* is approximate by using the potential stomatal conductance (*g_Stom_PAR_*) derived from Jarvis:

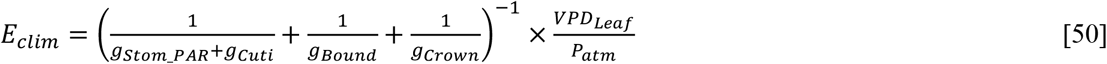 The next step is to determined how *E_Stom_* is regulated by the stomatal closure. When *E_Stom_* is not regulated then *E_Stom_* = *E_clim_* and *E_Leaf_*= *E_clim_* + *E_Cuti_*. Otherwise, *E_Stom_* is limited in order to maintain the target *P* (*e.g*. branch apoplasm) above the threshold potential reached when *E_Stom_* = 0. Let’s take the leaf apoplasmic potential as an example, the model first computes the equivalent conductance of the soil to leaf apoplasm pathway *K_Soil-to-L.apo_* and derive *E_Stom_* as:

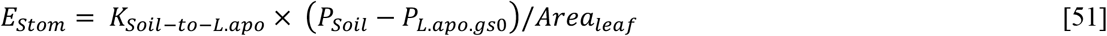 In these two first cases, once *E_Stom_* is known, *g_Stom_* can be easily back computed. It is important to note that as E_leaf_ = *E_Stom_*+*E_Cuti_*, hence transpiration remains lower-bounded by *E_cuti_* even when stomata are fully closed and *E_Stom_* = 0. For all organs, the elementary water movements *dq_organ_* from transpiration are finally computed as

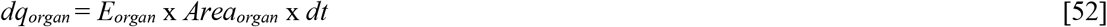

#### 2 Cavitation and redistribution of cavitated water

The percent loss of conductivity (PLC) is computed for the apoplasmic compartments of the different organs with a sigmoidal function. For a branch for instance at time *t*:

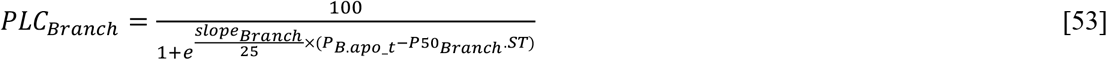

with *P50_Branch_*, the *P_B.apo_* corresponding to a PLC of 50 %, and *slope_Branch_* the slope of the curve at *P50_Branch_* and *ST* a factor accounting for the effect of temperature on water surface tension. By default, xylem refilling under negative pressure does not occur in ***SurEau***, and thus PLC can only increase under drought. The PLC increases in an organ as soon as water potential in the apoplasm continue to decrease below cavitation thresholds. When cavitation occurs, some apoplasmic water (*dQ_cavit_*) is released in the the system in proportion of the PLC:

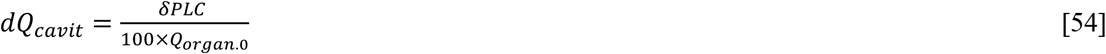

Where *δPLC* is the variation of cavitation between the current and previous time step. *dQ_cavit_* is distributed to associated symplasmic compartments. It is possible to activate a “refilling option”, which allows cavitated conduits to be refilled with surrounding symplasmic water, when xylem apoplasmic water potential increases above a user defined threshold (*P_refill_*).

#### 3 Physical properties dependent on temperature

Because one objective of ***SurEau*** was to predict plant water relations during heatwaves, we paid a special attention to the temperature dependence of the main physical properties of water solutions (see Cochard 2019 for more details). The reference values of the different parameters are taken at 20°C.

##### Fluidity

The dynamic fluidity *Fluid* of liquid water, the reciprocal of its viscosity, varies with temperature according to the empirical formula:

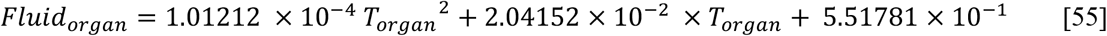

##### Surface tension of water

The surface tension of liquid water against air decreases with temperature according to this empirical formula:

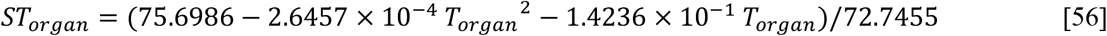

##### Osmotic potential temperature dependence

Following van’t Hoff relation, we define *OT* as:

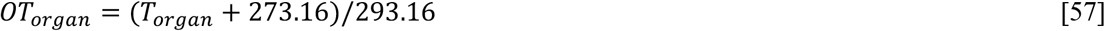

#### 4 Soil state variables (*P_soil_*, and *K_soil_* computed for each layer)

The hydraulic properties of the soil layers are defined by pedotransfer functions following van Genutchen (1980). Accordingly, the soil properties are characterized by 6 parameters (*θ_s_, θ_r_, α, n, K_sat_, l*). We describe the computation only for one layer.

The relative soil water content REW is the amount of water available between the water content *θ_fc_* at field capacity (*P_soil_*=0.033 MPa) and the residual water content *θ_r_*:

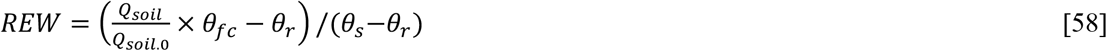

with *θ_s_* the soil water content at saturation. We assumed that soil water content cannot be higher than its value at field capacity.

The bulk soil water potential at *REW* is given by:

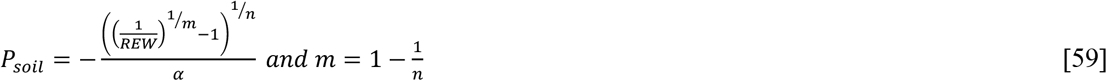

The soil hydraulic conductance at the interface with fine roots is computed as:

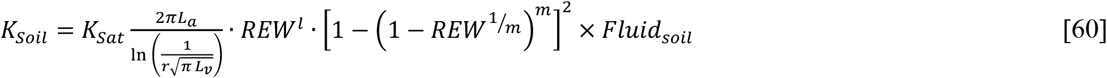

with *L_a_* and *L_v_* the root length per soil area and soil volume, respectively and *r* the radius of the roots.

The conductance of the interface between the soil and the root, *K_R.interf_* is empirically computed as a function of the root symplasmic shrinkage by dehydration:

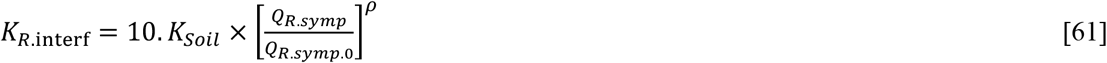

with ρ a parameter accentuating the isolation of the root system from the dry soil.

Finally, we compute the evapotranspiration from the soil surface *E_soil_* assuming a gaseous conductance *g_soil_* of the soil-atmosphere interface depending on the REW of the top soil layer as:

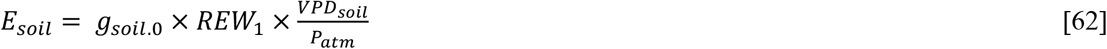

with *g_soil.0_* the conductance at soil saturation.

## III Results

To illustrate ***SurEau*** outputs, we modelled the hydraulic functioning of a 5m high young tree, with a diameter of 10cm, a leaf area of 10m^2^ and occupying a soil volume of 0.5m3. Tree physiological variables, climatic conditions, gaseous phase and soil parameters used in that simulation are given in Tables 3 to 6.

**Table 3:** Main physiological parameters of the plant organs and compartments.

| Organs | Parameters | Leaf | Branch | Trunk | Root |
| --- | --- | --- | --- | --- | --- |
| Symplasm | $\pi 0$ (MPa) | -2 | -2 | -2 | -2 |
| | $\epsilon$ (MPa<sup>-1</sup>) | 10 | 10 | 10 | 10 |
| | $K$ (mmol s<sup>-1</sup> MPa<sup>-1</sup>) | 25 | 6.1 | 0.77 | 8.33 |
| | $Q_0$ (mol) | 20.8 | 217.2 | 161.2 | 285.3 |
|  | Surface (m<sup>2</sup>) | 5 | 6.1 | 0.77 | 12.6 |
| Apoplasm | $P50$ (MPa) | -3 | -3 | -3 | -3 |
| | $Slope$ (% MPa<sup>-1</sup>) | 100 | 100 | 100 | 100 |
| | $K$ (mmol s<sup>-1</sup> MPa<sup>-1</sup>) | 50 | 98.8 | 414.3 | 222.3 |
| | $Q_0$ (mol) | 6.9 | 434.4 | 322.4 | 570.5 |
| | $C$ (mmol MPa<sup>-1</sup>) | 0.69 | 4.34 | 16.11 | 9.51 |

**Table 4:** Climatic conditions.

| Climatic conditions | $T_{air-min}$<br>°C | $T_{air-max}$<br>°C | $RH_{air-min}$<br>% | $RH_{air-max}$<br>% | $PAR$<br>μmol | $Wind\ speed$<br>ms<sup>-1</sup> | $VPD_{max}$<br>kPa |
| --- | --- | --- | --- | --- | --- | --- | --- |
| Values | 15 | 25 | 30 | 80 | 1500 | 1.0 | 2.0 |

**Table 5:** Main parameters for the flows in gaseous phase

| Gas phase Parameters | $g_{Stom\_max}$<br>mmol s<sup>-1</sup> m<sup>-2</sup> | $g_{Stom\_min}$<br>mmol s<sup>-1</sup> m<sup>-2</sup> | $g_{Cuti}$<br>mmol s<sup>-1</sup> m<sup>-2</sup> | $T_{phase}$<br>°C | $Q10a$ | $Q10b$ | $g_{Crown}$<br>mmol s<sup>-1</sup> m<sup>-2</sup> |
| --- | --- | --- | --- | --- | --- | --- | --- |
| Values | 200 | 10 | 2 | 35 | 1.2 | 4.8 | 200 |

**Table 6:** Soil parameters.

| Soil Parameters | volume<br>m<sup>3</sup> | Soil type | $\theta_s$ | $\theta_r$ | $\alpha$<br>cm<sup>-1</sup> | $n$ | $K_{sat}$<br>mmol/s/MPa | $l$ | $Q_{soil0}$<br>mol | $La$<br>m/m<sup>2</sup> | $Lv$<br>m/m<sup>3</sup> |
| --- | --- | --- | --- | --- | --- | --- | --- | --- | --- | --- | --- |
| Values | 1 | clay | 0.459 | 0.098 | 0.015 | 1.253 | 1.69 | 0.5 | 12000 | 206 | 412 |

For this test simulation, the plant is initiated in a soil at its field capacity and allowed to dehydrate gradually until being completely dry (water inputs from precipitation are assumed to be equal to zero). *g_Stom_* was here modelled with P_*gs10*_ = −1.5 MPa and P_*gs90*_ = −2MPa (equation [49]).

At the beginning of the simulation, the daily variations of the different physiological variables of the plant follow the daily climatic variations, when the soil is still wall-watered (Figure 3). The stomatal conductance is mainly light-controlled, but the dynamics of transpiration and water potentials are slightly delayed with respect to this conductance because the VPD peak is reached 2 hours after solar noon. The model therefore captures well these complex responses of stomata and transpiration to the different climatic variables. When dehydration is maintained, the stomata progressively close according to the intensity of foliar water stress (Figure 4). After about thirty days, the stomata are permanently closed and transpiration is limited to cuticular losses which gradually accentuate the hydric stress of the plant. At this stage, cavitation events begin in the apoplasm of the various organs, decreasing the amount of water stored in the vessels. When the embolism rate of the leaf apoplasm reaches 100% (after 80 days), the leaf water potential drops abruptly (to reach the water potential of the air) as leaves lose their symplasmic water stock. The hydraulic failure of the leaf xylem tissue causes desiccation. Later (day 100), the same phenomenon occurs for the branches, and finally for the trunk.

**Figure 3:**
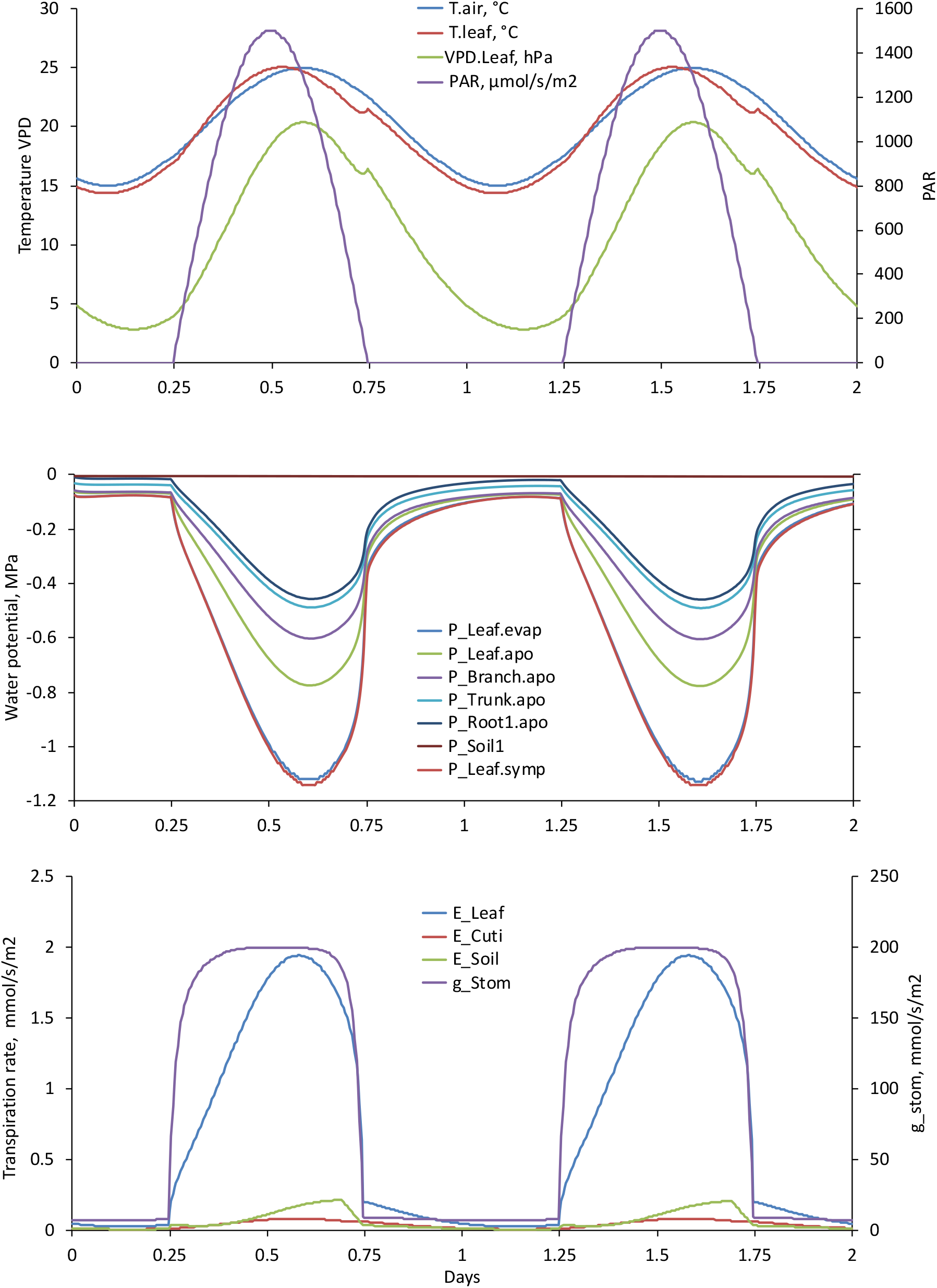
Daily variations of key physiological variables simulated with *SurEau*.c. Two consecutive days are simulated for a well-hydrated plant.

**Figure 4:**
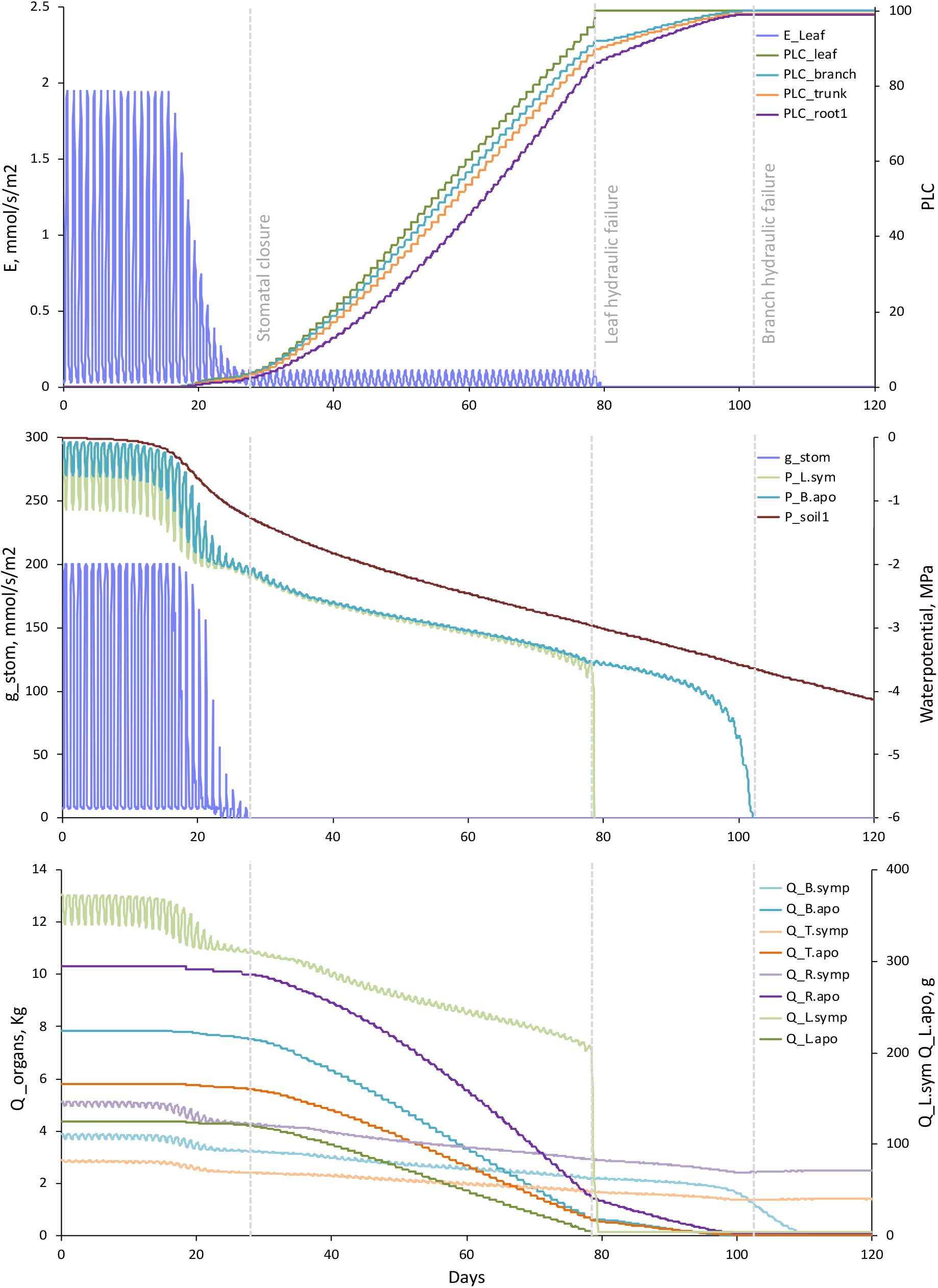
Simulation of the effect of an extreme water stress with *SurEau*.c. Simulation of different key tree variables is shown for 120 consecutive days. The tree is placed in well-watered soil at t=0 and allowed to dehydrate until complete desiccation. At t=28 days stomata are closed. At t=80 days PLC reaches 100% in the leaf apoplasm provoking leaf desiccation. and t= 102 days hydraulic failure occurs in the branch apoplasm.

## IV Discussion

The ***SurEau*** model uses classic bioclimatic and hydraulic formalisms to account for gas exchanges and the water relations of a plant. This modeling is based on water mass conservation and on a parameterization of hydraulic and hydric properties and apoplasmic and symplasmic properties of organs (roots, trunk, branches, leaves). This idealization offers a good compromise between complexity (computation time) and reliability of the representation of the processes we are interested in. The number of parameters in the model remains quite high, but an accurate parameterization of the model remains possible. A number of experimental devices-not described here-have been developed and are still development to estimate the parameters that are the most difficult to measure. For example, a fractal representation of the aerial and root parts allows to estimate the volumes and exchange surfaces of these organs.

The main limitation of our model is currently its computational cost, an inevitable consequence of the need to model dynamic processes with a very short characteristic time imposed by the CFL constraint. For example, the simulation shown in figure 4 required a calculation time of 2 minutes on a PC with a powerful processor (AMD 2970WX). This execution time is not a constraint to simulate an isolated tree, but can be critical to simulate long climatic series on network grids for instance. To meet this need, other numerical integration techniques of ***SurEau*** (steady-state approach, hyperbolization, Durdorf-Frankel scheme) are under development and testing to speed up the code.

The main contribution of ***SurEau***.c to existing models is the detailed responses to extreme water stresses of the hydric, hydraulic and gaseous functioning of the plant. The model can capture the effect of water stress on the induction of cavitation, and thus of a xylem hydraulic failure on the desiccation of organs supplied by this tissue. A key advantage is the possibility to track the water storage in the plant, which can be expressed per unit volume, area, or dry mass as it relates to fire danger and remote sensing indicators. ***SurEau*** therefore appears to be a particularly suitable tool for better understanding the effects of extreme water stress on plant survival. This type of model is expected to better predict and anticipate the future effects of climate change on the survival of plant species.

## VI Acknowledgments

HC received found from the ANR projects 16-IDEX-0001 and 18-CE20-0005.

